# Genetic Association of *CSN2* Variant 6:85451298C>A with A1/A2 Milk Phenotype in Pakistani Cows

**DOI:** 10.1101/2022.08.25.505361

**Authors:** Sibghat Ullah, Zubair Ahmed, Asma Karim, Rashid Saif

## Abstract

The cow is a very important mammal from the bovine family renowned for providing milk, meat and hide around the globe. β-casein is one of the principal proteins found in cow’s milk encoded by the *CSN2* gene with 12-13 reported genetic variants present on Chr.6 in the cow’s genome. The current study investigated A1/A2 associated variant g.6:85451298C>A (rs43703011) in exon 7^th^ according to the GenBank ID: M55158.1 position 8101C>A (c.245C>A), which is responsible to change the 67^th^ amino acid of the β-casein polypeptide from proline (CCT, A2) to histidine (CAT, A1). The milk containing A2 protein variant is good while A1 milk has some negative effects on human health due to the production of β-casmomorphin-7. Genetic screening of the subject variant was conducted using ARMS-PCR on 48 native and exotic cows from Pakistan. Our initial results showed that 37.5% of the population is homozygous wild-type (C/C), 56.25% is heterozygous (C/A) and the remaining 6.25% is homozygous mutant-type (A/A). Additionally, the Chi-square association test was applied using the PLINK data analysis toolset which presented a significant *p-*value of 2.811×10^−3^ and OR of 3.8 with an alternative allele frequency of 0.2 and 0.5 in native and exotic populations respectively. Hence, there is significant variability in A1/A2 genotype which may be addressed by adopting selective breeding programs in Pakistan.

## Introduction

Milk is considered one of nature’s complete food for infants and as well as for adults because it contains all the required and essential nutrients [1]. Cow milk is basically 85-87% water, 4.8-4.9% lactose sugar (carbohydrates), 3.2-3.8% proteins, 3.7-4.4% fats and 0.7-0.8% minerals (Ash). Casein is one of the major milk proteins found in 4 subtypes: (alpha) αs1-casein, (alpha) αs2-casein, (beta) β-casein, and (kappa) κ-casein. β-casein is one of the significant proteins which is a primary source of essential amino acids for a suckling newborn. This protein constitutes 35% of total milk casein made up of 209 amino acids long polypeptide with molecular weight of 23,983kDa [2].

The 10,338 bp long bovine *CSN2* gene located on Chr.6, encode β-casein protein. Almost 12 variants of this gene have been reported so far i.e. (A1, A2, A3, B, C, D, E, F, G, H1, H2 and I) which code for different β-casein proteins in milk [3]. A1/A2 are most studied and globally renowned variants due to their significant association with A1/A2 milk phenotype. This variation occurs due to a single nucleotide polymorphism (SNP) which results in replacement of wild-type amino acid proline into mutant-type histidine at 67^th^ position in β-casein polypeptide. According to the literature, domestication, natural selection and random breeding results in a point mutation in *CSN2* gene which changed wild-type A2 allele into its mutant-type A1[4].

Nowadays, presence of A1/A2 β-casein in milk describes its quality. As the mutant type (A1) possess histidine residue, this result in a very weak bond between 66^th^ and 67^th^ position in its polypeptide. Upon proteolytic digestion the bond breaks and release a seven amino acid (Tyr-Pro-Phe-Pro-Gly-Pro-Ile) containing bioactive peptide known as β-casomorphin-7 (BCM-7) but A2 β-casein does not releases it [5, 6]. BCM7 is believed to cause human health problems as it can potentially affect number of opioid receptors in the endocrine, nervous, and immune system. It is identified as oxidizing low dietary lipoproteins (LDL) which result in hardening and thickening of blood arteries. β-casein A1 milk is also associated with type-1 diabetes, sudden infant death syndrome, chronic constipation in infants, bloating and irritable bowel syndrome due to delayed gastrointestinal activity [7-9].

The following research is designed to investigate genetic association of *CSN2* gene with A1/A2 milk phenotype in Pakistani cow population. *CSN2* A1/A2 allele is located on Chr.6, accession ID NC_037333.1 in assembly ARS-UCD1.3(GCF_002263795.2) in which variation at region 6:85451298C>A (rs43703011) in exon 7 according to GenBank accession ID: M55158.1, transcript position 8101 with CDS c.245C>A, protein variant AAA30431.1 change the 67^th^ amino acid of β-casein polypeptide from proline (CCT, A2) to histidine (CAT, A1) [10].

## Materials and Methods

### Sample collection and DNA extraction

Native and exotic cow breeds were selected to study association of *CSN2* locus (6:85451298C>A) with A1/A2 milk phenotype in Pakistani cows. Genomic DNA was extracted from cow blood. Forty-eight blood samples were collected from the abattoir “Punjab Agriculture & Meat Company” (PAMCO), Multan Rd, Lahore, Pakistan. Blood was taken into K3-EDTA vacutainers and stored at 4^°^C till further use. Organic method was used to extract DNA from the samples.

### ARMS primer designing

Based on GenBank accession ID:M55158.1 with complete CDS, ARMS-PCR primers were designed using Primer3 (https://primer3.ut.ee/) and OligoCalc (http://biotools.nubic.northwestern.edu/OligoCalc.html) software. Five primers were designed, three of which were (i) reverse normal (N) ARMS (ii) reverse mutant (M) ARMS primers from the 3’-end to amplify wild-type and mutant variants of the targeted sequence, along with (iii) a forward common primer. To improve the specificity and effectiveness, a secondary mismatch was also added at the fourth nucleotide position from the 3’ end of reverse wild/mutant ARMS primers. Additionally, two more primers (forward and reverse) were designed to amplify a region near the targeted sequence that serve as internal control (IC).

### PCR amplification

To amplify wild and mutant-type variants SimpliAmp thermal cycler (Applied Biosystems) was used. Normal and mutant ARMS reverse primer along with common forward primer were analyzed in two separate PCR reactions for each sample. Internal control (IC) primers were also added in each tube at the same time to amplify nearby targeted locus. One µL of extracted genomic DNA, 1µL of each primer (forward common primer, N or M-reverse primer, forward and reverse IC primers), one unit of *Taq* polymerase, 2.5 mM of MgCl_2_, 2.5 mM of dNTPs, 1x buffer, and PCR-grade water were pipetted to make a total reaction volume of 25µL. In the PCR protocol, initial denaturation at 95°C for 5 minutes, following by 30 cycles of denaturation at 95°C for 30 sec, annealing at 60°C for 30sec, and extension at 72°C for 30sec with final extension lasted for 10 minutes at 72°C.

### Statistical association

The genotypic and allelic diversity of g.6:85451298C>A locus in our sampled cows were analyzed followed by chi-square association test using PLINK data analysis toolset for genotypic and allelic frequencies along with their *p*-value and odds ratio (OR).

## Results

### Variant genotyping

Genotypic variability of g.6:85451298C>A was analyzed in 24 native and 24 exotic (total 48) cow samples from Pakistan. Significant genotypic variability of this variant was shown in ARMS-PCR results which associates with the production of A1/A2 β-casein protein in cow’s milk. Out of total 48 cows, 18 had wild (C/C) genotype associated with A2 protein, 27 cows had heterozygous C/A genotype associated with both A1/A2 protein and only 3 showed mutant (A/A) genotype associated with A1 protein. Some pictures of sampled cows alongside results are shown in figure 2.

**Figure 1:**
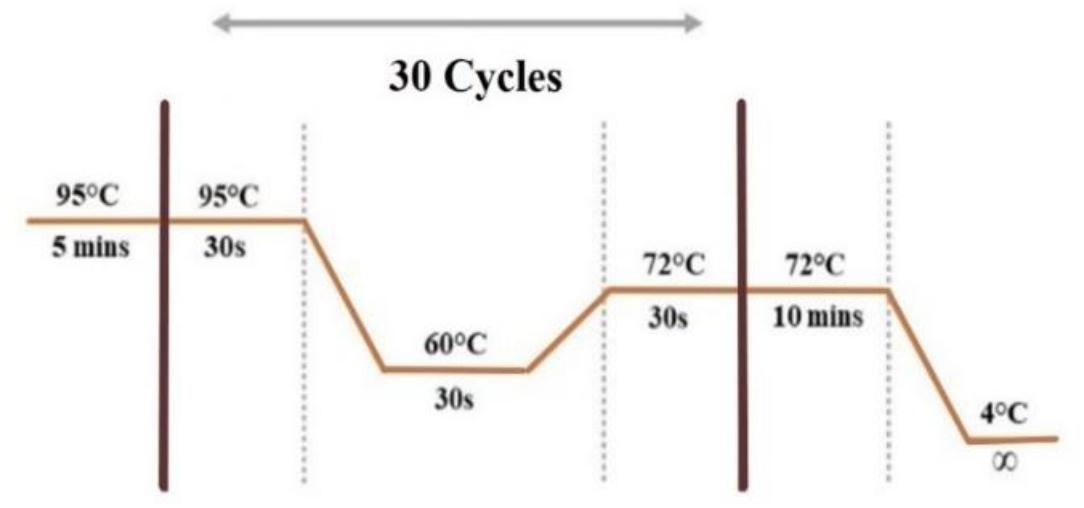
Thermal Cyclic Conditions of ARMS-PCR

**Figure 2:**
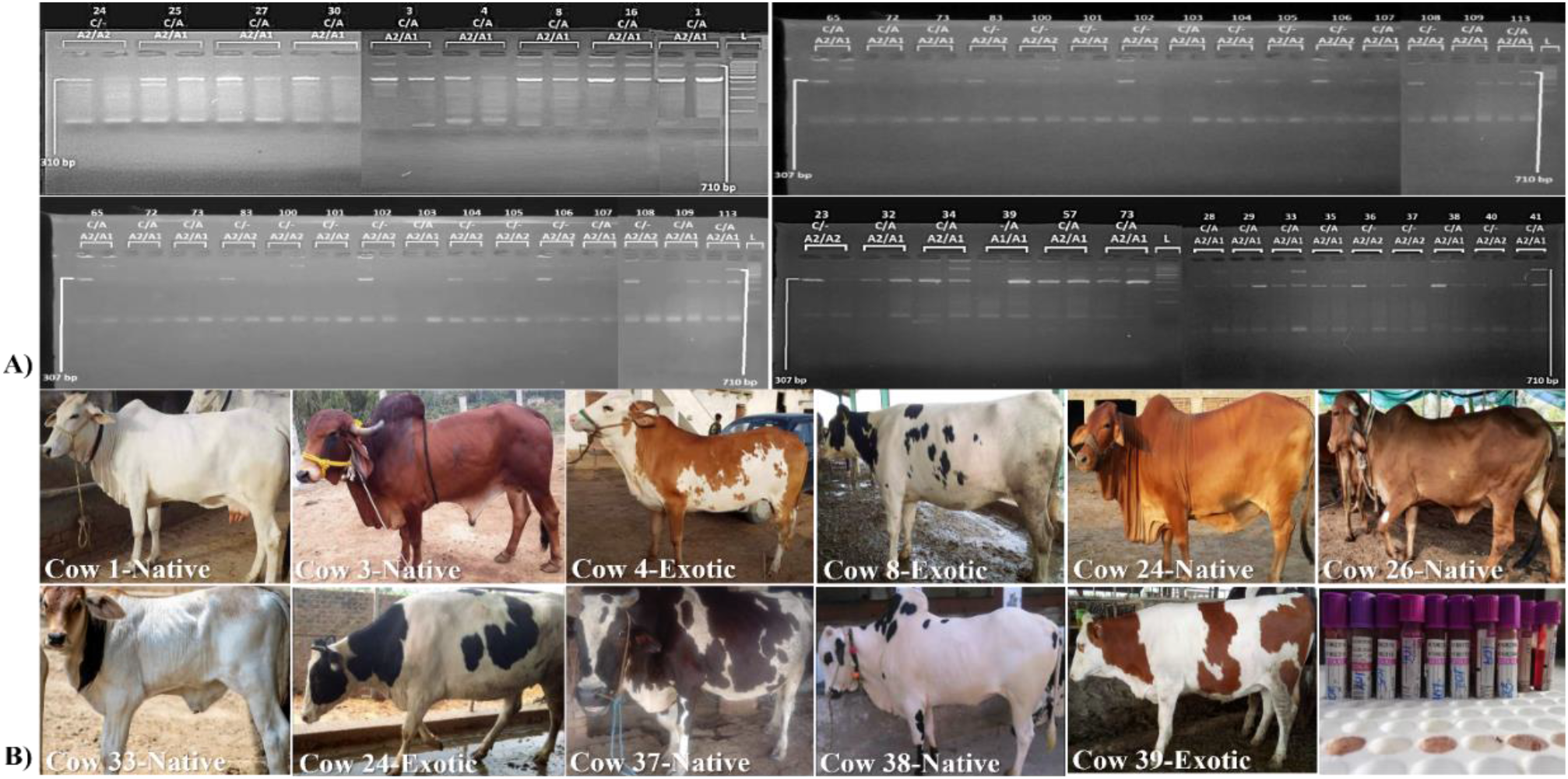
**A)** ARMS-PCR amplification of targeted variant **(B)** Photos of some sampled animals

### Association analysis

Chi-square test was used to compare native and exotic population by determining their minor allele frequencies (MAF). These frequency distributions demonstrated that both native and exotic Pakistani cows often had the heterozygous (C/A) genotype codes for both A1/A2 β-casein variant in milk. Screening of our samples showed, 37.5% of the cow population was homozygous wild (C/C), 56.25 percent was heterozygous (C/A), and the remaining 6.25 percent was homozygous mutant (A/A). Additionally, the findings of ARMS-PCR genotyping revealed that “C” allele was more prevalent than the “A” allele. A significant *p*-value of 2.811×10^−3^ was computed using PLINK data analysis toolset (Table 2).

**Table 1:**
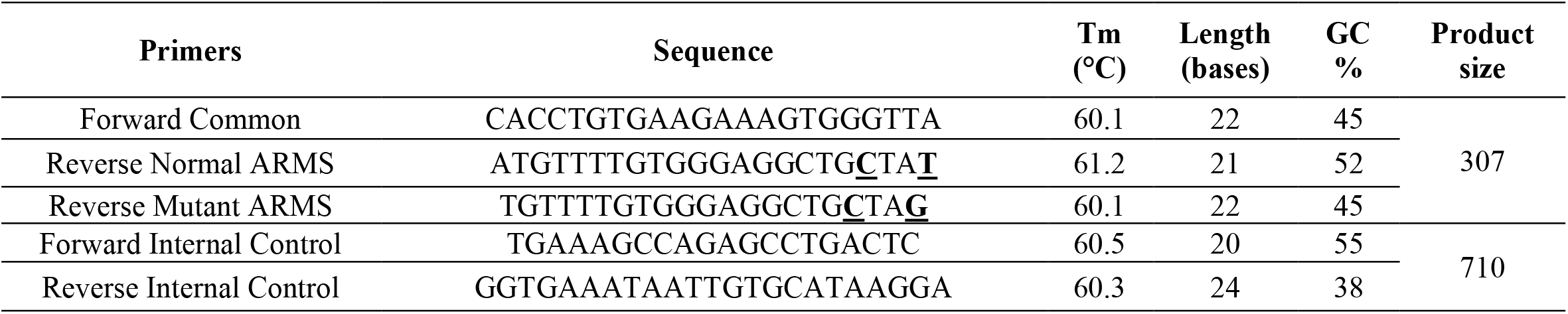
ARMS-PCR Primers Sequences

**Table 2:** Association Analysis of Subject Variant in Cow Population

| # of samples | Genomic position | cDNA variant<br>M55158.1 | Protein variant<br>AAA30431.1 | Genotypic information | | | Alt. allele frequency | | $p$-Value (OR) |
| --- | --- | --- | --- | --- | --- | --- | --- | --- | --- |
|  |  |  |  | Homo-wild (CC/%) | Hetero- (CA/%) | Homo-mutant (AA/%) | Exotic | Native |  |
| 48 | g.6:85451298 | c.245C>A | p. P67H | 18/37 | 27/56 | 3/6 | 0.5 | 0.2 | $2.811 \times 10^{-3}$ (3.8) |

## Discussion

Bovine *CSN2* gene is associated with A1/A2 milk phenotype having 12-13 different β-casein variants reported in the literature [3]. The locus 6:85451298C>A which particularly got attention of researchers around the globe to explain the genetics of A1/A2 milk in different cows breeds [10]. β-casein A1 variant is supposed to release a biologically active peptide β-casomorphin-7 (BCM-7) which is believed to have negative effects on human health e.g., blotting, type-1 diabetes, sudden infant death syndrome (SIDS), bowel syndrome, delayed gastrointestinal activity and heart diseases [7-9].

Out of total 48 screened samples, 24 native and 24 exotic cows were selected to investigate the genetic association of *CSN2* gene with A1/A2 milk phenotype along with the statistical association analysis revealed genotypic and allelic frequencies of our sampled population. Current results showed overall 37.5% of the cow population was homozygous wild (C/C), 56.25% was heterozygous (C/A), 6.25% was homozygous mutant (A/A) and significant *p-*value of 2.811×10^−3^ with alternative allele frequency of 0.2 and 0.5 in native and exotic cows respectively. As the alternative allele frequency in native is lesser in our population as compare to exotic cows that is in conformity to alternative allele frequency of native animals of the regional countries like India and Bangladesh [11, 12]. While alternative allele frequency in Pakistani exotic animals is not much higher, as expected due to the random and unplanned breeding which is also introducing A1 allele in our native cow population just to enhance the milk production without considering the A1/A2 milk quality unlike the western and Australian cows.

## Conclusion

Genetic variations in *CSN2* gene marker (6:85451298C>A) is statistically significant with A1/A2 milk phenotype in Pakistani native cow population with moderate induction of A2 allele in exotic animals due to random breeding, but at the same time, our local cows are also getting A1 allele which is not appreciated for local farmers. As a nutshell, current study suggests the screening of the cattle herds in Pakistan to maintain the genetic potential of indigenous and exotic animal as their exclusive genetic resources.

## Ethical Statement

For this research, ethical guidelines and permission are omitted. There was no special need for approval from an animal ethics committee. Sampled cows belong to the abattoir “Punjab Agriculture & Meat Company” (PAMCO), Lahore Pakistan which were approached through proper channels via permission letter provided as supplementary material in this manuscript.

## Acknowledgement

Authors are obliged to the staff and employees of PAMCO for their support to provide blood samples and accomplish this research work.

## Authors Contribution

Sibghat Ullah (SU) and Zubair Ahmed (ZA) has equal contribution as first author, collected samples, performed wet-lab and wrote the initial draft; Rashid Saif (RS) and Asma Karim (AK) designed the project, trouble shooting in wet & dry-labs, arranged samples, applied association testing, editing and proof reading of the manuscript.

---

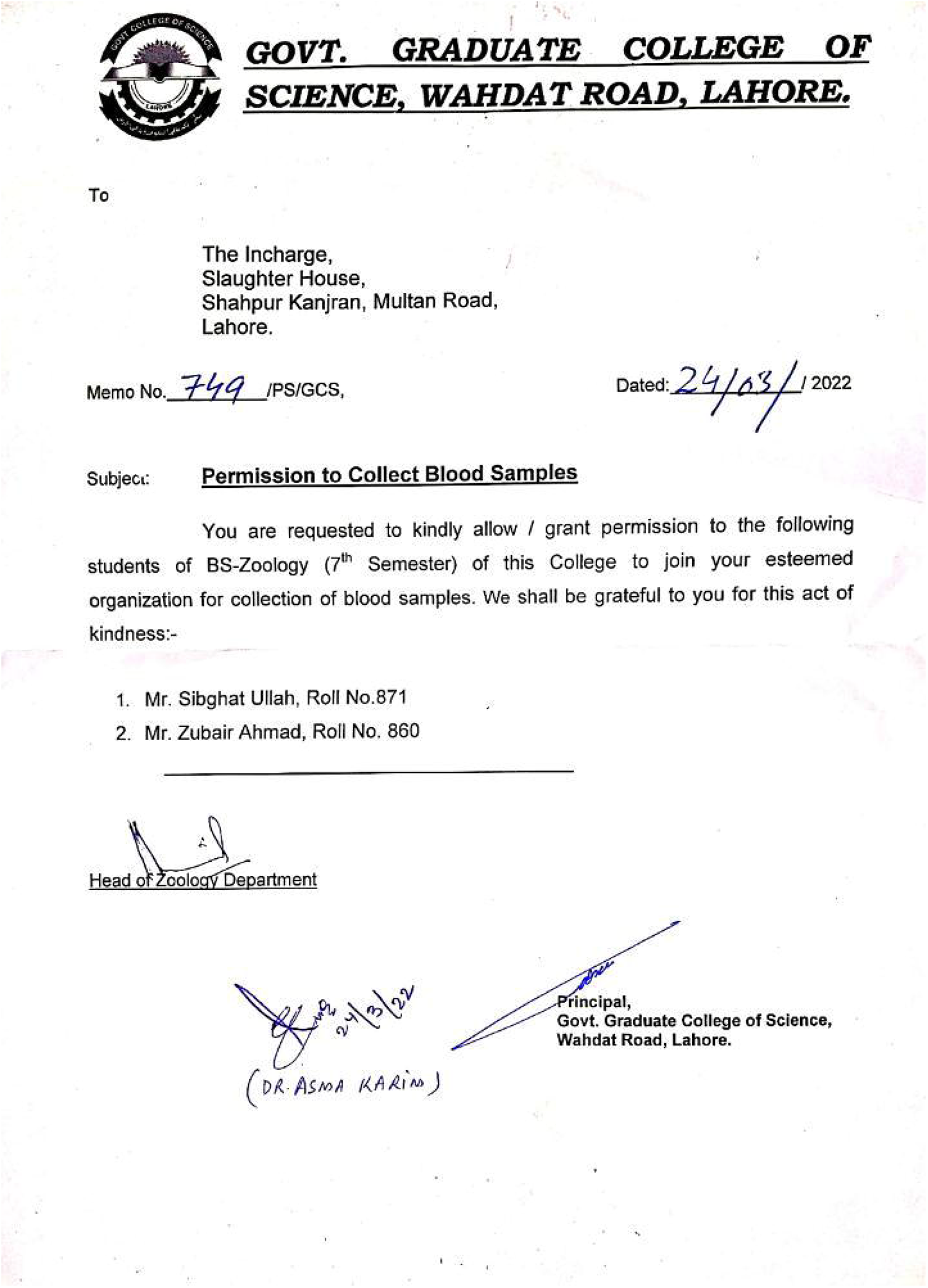

